# Tryptophan metabolites as biomarkers to predict the severity and prognosis of acute ischemic stroke patients

**DOI:** 10.1101/2025.02.24.640001

**Authors:** Chuzheng Pan, Feng Chen, Yan Yan, Haiwen Li, Chengfeng Qiu

**Author notes:** Chuzheng Pan, Feng Chen and Yan Yan contributed equally to this work.

## Abstract

**Background:** A growing body of evidence indicates alterations in metabolite levels and enzyme activities associated with the conversion of tryptophan (TRP) throughout the course of cerebral ischemia. In this study we aim to explore the potential relationship between TRP metabolism and clinical prognosis in acute ischemic stroke (AIS) patients of mainland China.

**Methods:** Blood samples were obtained from a cohort of 304 patients diagnosed with AIS. The concentrations of ten TRP metabolites were quantified utilizing liquid chromatography-tandem mass spectrometry (LC-MS/MS). Stroke severity was evaluated upon admission using the National Institutes of Health Stroke Scale (NIHSS). A poor functional outcome was defined as modified Rankin scale (mRS) > 3, whereas a good functional outcome was defined by mRS ≤ 3 at 3 months post-stroke. LASSO regression and random forest algorithms were then employed to identify key TRP metabolism parameters associated with prognosis.

**Results:** Following the optimization of variable selection through Lasso regression, a prognostic risk model with 7-factors related to AIS was constructed, yielding an AUC of 0.917. Subsequently, a random forest analysis was conducted to establish an 11-factor prognostic risk model, which demonstrated an enhanced AUC of 1.000. Ultimately, three robust parameters related to TRP metabolism were identified. Multivariable logistic regression analysis, adjusted for covariates, revealed that TRP (odds ratio [OR] = 0.46, 95% confidence interval [CI]: 0.26 - 0.76, *p* = 0.004), the kynurenine (KYN)/TRP ratio (OR = 2.06, 95% CI: 1.23 - 3.60, *p* = 0.008), and the kynurenic acid (KYNA)/TRP ratio (OR = 2.15, 95% CI: 1.23 - 4.12, *p* = 0.014) were independently associated with poor functional prognosis.

**Conclusions:** The results of this study indicate that TRP metabolism is associated with the severity and prognosis of AIS. The TRP, KYN/TRP ratio and KYNA/TRP ratio may serve as potential biomarkers for 3-month prognostic evaluation.

## 1. Introduction

Stroke is the second leading cause of death worldwide and is also among the leading causes of long-term adult disability [1]. Despite the striking improvement of acute stroke management during past decades involving thrombolytic treatment, thrombectomy and stroke inpatient units [2], stroke related mortality and morbidity remain to be a major health concerns. The often severe and detrimental consequences of stroke impose a substantial burden due to physical disability and associated healthcare costs [3]. Therefore, prediction of stroke outcome in an early stage is pivotal in order to implement early rehabilitation when necessary. Although age, initial clinical severity and brain imaging are established clinical predictors for long-term outcome [4, 5], blood biomarkers can be a complementary tool for diagnosis [6], predicting prognosis [7, 8] and therapeutic monitoring of novel treatments in ischemic stroke [9].

The majority of stroke cases are ischemic, primarily resulting from superimposed thrombosis on underlying atherosclerotic plaques within the cerebral vasculature[1]. The pathogenesis of atherosclerosis closely connects activation of proinflammatory signaling pathways, expression of cytokines/chemokines and production of reactive species [10, 11]. Tryptophan (TRP) is an important essential amino acid that plays an indisputable role in various physiological processes, notably neuronal function and immune responses [12]. The metabolism of TRP in the human body proceeds through three different pathways, including the kynurenine (KYN), serotonin (5-HT) and indole pathways. Furthermore, other biologically active components, such as 5-HT, melatonin, and niacin are by-products of TRP pathways [12, 13]. A growing body of evidence illustrates a important role for the TRP catabolism in ischemic stroke[14–17]. The increased TRP catabolism is initiated before or immediately after stroke and remains altered during the acute phase [18], indicating its potential as a promising biomarker for prognosis of AIS.

Considering the variations in geographical regions, ethnic groups and dietary habits, few studies have explored TRP catabolism in all three pathways in mainland China. We therefore investigated the metabolites of TRP in plasma of acute ischemic stroke (AIS) patients and their relation with initial stroke severity and long-term outcome.

## 2. Materials and methods

### 2.1 Study participants

This study included consecutive patients with AIS between March 2024 and September 2024. We recruited 304 patients with acute ischemic stroke (AIS) confirmed by magnetic resonance imaging (MRI) or computed tomography (CT) of the brain within 14 days of symptom onset. The other inclusion criterion was age ≥ 18 years. We excluded patients with disabilities (Modified Rankin Scale score ≥ 2) before stroke onset and those who underwent intravenous thrombolysis or mechanical thrombectomy. This study was approved by the ethics committee of the Hunan University of Medicine General Hospital.

### 2.2 Clinical Assessments

We assessed demographic characteristics and comorbidities, including age, sex, body mass index (BMI), blood pressure, hypertension, diabetes mellitus, coronary heart disease (CHD), atrial fibrillation, history of stroke, smoking and moderate or heavy drinking. Kidney function, blood routine examination, fasting blood glucose (FBG), HbAlc, coagulation function, creatine kinase, C-reactive protein (CRP), thyroid function, erythrocyte sedimentation rate (ESR) and serum lipids were determined from each patient after admission to hospital. The National Institutes of Health Stroke Scale (NIHSS) was used to assess the severity of stroke [19]. Laboratory examinations regarding the glucose profile, lipid profile and HbAlc were performed after 12 h of overnight fasting.

Clinical information ragarding patient outcomes after discharge was prospectively collected during routine clinic visits or via telephone interviews with patients or their caregivers, conducted 3 months after the qualifying event. The primary outcome of this study was poor functional outcome, which was assessed using the modified Rankin Scale (mRS). A poor functional outcome was defined as mRS > 3, whereas a good functional outcome was defined by mRS ≤ 3 at 3 months post-stroke.

### 2.3 The measurements of TRP metabolites

#### 2.3.1 Chemicals and reagents

TRP (99.3%), KYN (99.9%), N-formylkynurenine (NFK) (97.1%), 3-hydroxyanthranilic acid (3-HAA) (99.3%), 5-HT (95.9%), 5-hydroxyindoleacetic acid (5-HIAA) (97.6%), 5-hydroxytryptamine (5-HTP) (99.3%) and indole-3-acetic acid (IAA) (99.7%) were purchased from Shanhai Aladdin Biochemical Technology Co., Ltd (Shanhai, China). kynurenic acid (KYNA) (100%) and 3-HKYN (98.9%) were purchased from Sigma-Aldrich (St Louis, Missouri, USA). Chemicals purity were provided in brackets for each standards molecules. Acetonitrile and methanol (HPLC-grade) were purchased from Merck (Darmstadt, Canada). Formic acid (AR-grade) was purchased from Shanhai Aladdin Biochemical Technology Co., Ltd (Shanhai, China). Phosphate-buffered saline (PBS) was purchased from Beijing Dingguo Biochemical Co., Ltd (Beijing, China). Ultrapure water was prepared using a GenePure XCAD plus system procured from ThermoFisher Scientific (Waltham, USA).

#### 2.3.2 Instrumentation and liquid chromatography-tandem mass spectrometry (LC-MS/MS) conditions

Chromatography was performed on an ACQUITY UPLC I-Class system, equipped with an ACQUITY UPLC I-Class BSM (Quaternary pump), an ACQUITY UPLC I-Class SM-FTN (Autosampler) and an ACQUITY UPLC I-Class CH-A (Column oven) (Waters, Zellik, Belgium). Samples were thermostated at 10℃ and the separation was performed at 40℃ using a Boltimate® EXT-C18 Core-Shell column (2.1 mm×100 mm, Particle size: 2.7 μm, Welch Materials, Inc., China). The mobile phase consisted of solvent A (deionised water with 0.5% formic acid) and solvent B (acetonitrile with 0.1% formic acid). The gradient elution program was as follow:mobile phase B maintained at 7% (0.00-1.00 min), from 15% to 40% (1.01-3.50 min), maintained at 95% (3.51-4.80 min), maintained at 6% (4.80-5.50 min). The gradient program uses a flow rate of 0.25 mL/min for 1.01-3.60 minutes, and a flow rate of 0.40 mL/min for the other times periods. The valve was diverted to MS between 0.40-3.20 minutes. The injection volume was 8 μL.

Quantitative of TRP and its metabolites was achieved using a Xevo TQD MS/MS (Waters, Zellik, Belgium). Data acquisition was performed on Masslynx® 4.2 software. Detection of the ions was performed in the multiple reaction monitoring (MRM) mode at unit mass resolution. Electrospray ionization (ESI) was operated in positive mode. Critical MS parameters such as transition, collision energy and cone were carefully optimized to obtain the best possible sensitivity. For each analyte, the MS conditions were determined via direct infusion of individual standard solution. Source temperature, desolvation temperature, desolvation source gas flow, cone gas flow and capillary voltage was set at 150 ℃, 600 ℃, 1000 L/Hr, 20 L/Hr and 1.00 kV. The optimized parameters are shown in Supplement Table 1.

#### 2.3.3 Preparation of standard solutions, calibration standards and QCs

Based on the solubility of TRP and its metabolites, stock solutions were prepared as follows: TRP (5.00 mg/mL), NFK (1.00 mg/mL), and KYN (1.00 mg/mL) were made using a 50/50 (v/v) water/methanol mixture. Stock solutions of 5-HT (1.00 mg/mL), 5-HIAA (1.00 mg/mL), IAA (1.00 mg/mL), and 5-HTP (1.00 mg/mL) were prepared using pure methanol. Stock solutions for 3-HAA (1.00 mg/mL) and KYNA (1.00 mg/mL) were prepared using 100% DMSO, while the stock solution of 3-HKYN (1.00 mg/mL) was made with DMSO containing 0.1% formic acid. To address potential interference among TRP and its nine metabolites during the synthesis process, we created two working solutions: Working Solution 1 (containing NFK, 3-HKYN, and 5-HTP) and Working Solution 2 (containing TRP, KYN, 3-HAA, KYNA, 5-HIAA, IAA, and 5-HT). Both working solutions were stepwise diluted with 50% methanol in water to generate standard curve working solutions at eight concentration levels and quality control working solutions at four concentration levels (including LLOQ QC, LQC, MQC, and HQC). All prepared stock and working solutions were stored at -80°C prior to analysis.

Given that TRP and its metabolites are endogenous substances and blank serum is difficult to obtain, Calibration standards at eight levels were prepared by spiking the mixed working solutions in 1M PBS at the volume ratio of 1:19 (1.50, 3.00, 7.50, 15.0, 30.0, 60.0, 120, 150 ng/mL for NFK; 1.00, 2.00, 5.00, 10.0, 20.0, 40.0, 80.0, 100 ng/mL for 3-HKYN and 5-HTP; 20.0, 40.0, 100, 200, 400, 800, 1600, 2000 ng/mL for KYN; 1.00, 2.00, 5.00, 10.0, 20.0, 40.0, 150, 300 ng/mL for 5-HT;1.00, 2.00, 5.00, 10.0, 20.0, 40.0, 80.0, 100 ng/mL for 3-HAA and KYNA; 200, 400, 1000, 2000, 4000, 8000, 16000, 25000 ng/mL for Trp; 3.00, 6.00, 15.0, 30.0, 60.0, 120, 240, 300 ng/mL for 5-HIAA; 10.0, 20.0, 50.0, 100, 200, 400, 800, 1000 ng/mL for IAA). In contrast, the quality control (QC) samples at low (LQC), medium (MQC) and high (HQC) levels were prepared by spiking the mixed working solutions in 1M PBS at the volume ratio of 1:19 (4.50, 45.0, 90.0 ng/mL for NFK; 3.00, 30.0, 60.0 ng/mL for 3-HKYN and 5-HTP; 80.0, 600, 1400 ng/mL for KYN; 4.00, 30.0, 120 ng/mL for 5-HT; 4.00, 30.0, 70.0 ng/mL for 3-HAA and KYNA; 800, 6000, 14000 ng/mL for Trp; 12.0, 90.0, 210 ng/mL for 5-HIAA; 40.0, 300, 700 ng/mL for IAA).

#### 2.3.4 Serum sample preparation

Blood sampels were collected from the patients after admission, following an overnight fast. The samples were processed using a simple protein precipitation method. A total of 100 μL of the sample (including standard curve samples, quality control samples, and study samples) was transferred to a 1.5 mL Eppendorf tube. Then, 100 μL of acetonitrile:methanol (7:3, v/v) precipitant was added, and the mixture was vortexed for another 5 minutes. The samples were then centrifuged at 13,000 rpm for 15 minutes at 4°C. Subsequently, 60.0 μL of the supernatant was transferred to a 1.5 mL Eppendorf tube containing 300 μL of pure water and vortexed for 5 minutes. The prepared samples were then transferred to a 96-well plate for analysis using the LC-MS/MS system.

### 2.4 Statistical analysis

Continuous variables were reported as medians with interquartile ranges (IQR), and categorical variables were expressed as frequencies and percentages. The Mann-Whitney U-test was employed for the analysis of continuous variables, whereas the Chi-square test was utilized for categorical variables. The correlation analysis between TRP metabolic parameters and disease severity was tested using Spearman correlation analysis. To identify the most robust variables, the least absolute shrinkage and selection operator (LASSO) regression and random forest methods were applied. Model accuracy was evaluated using the receiver operating characteristic (ROC) curve. Common features were identified using a Venn diagram analysis. Furthermore, multiple logistic regression analysis was conducted to examine the associations between TRP metabolic parameters and the risk of a poor prognosis at 90 days in patients with cerebral infarction. A two-tailed *p* value <0.05 was considered statistically significant. All statistical analyses were performed using the R software (version 3.4.2).

## 3. Results

### 3.1 Patient characteristics and TRP parameters

The baseline characteristics of the individuals in the current analysis are summarized in Table 1. In total, TRP metabolite measurements were performed in 304 patients with AIS. Of the all the patients, 49 (16.12%) patients had mRS scores > 3 (poor prognosis, Ppoor group), and 255 (83.88%) patients had mRS scores ≤ 3 (good prognosis, Pgood group). Univariate analysis revealed that individuals in the Ppoor group exhibited significantly higher values for age, prevalence of atrial fibrillation, white blood cell count (WBC), CRP, blood urea nitrogen (BUN), prothrombin time (PT), international normalized ratio (INR), free thyroxine (FT), NIHSS scores compared to those in the Pgood group (*p* < 0.05). Conversely, triglycerides (TG), prothrombin activity (PTA), and free triiodothyronine (FT3) were significantly lower in the Ppoor group than in the Pgood group (*p* < 0.05).

**Table 1.** Baseline characteristics of the patients with cerebral infarction in the poor and good prognosis groups.

| Variables | Pgood(n=255) | Ppoor(n=49) | $p$ |
| --- | --- | --- | --- |
| **Demographic** |  |  |  |
| Age, Median(IQR), yr | 63.00 (56.00, 71.50) | 73.00 (64.00, 78.00) | $< 0.001$ |
| BMI, Median(IQR), kg/m<sup>2</sup> | 23.90 (22.25, 26.25) | 23.30 (20.40, 25.80) | 0.076 |
| Male Gender, n(%) | 170 (66.67) | 30 (61.22) | 0.462 |
| SBP, Median(IQR), mmHg | 146.00 (133.50, 159.00) | 149.00 (137.00, 162.00) | 0.255 |
| DBP, Median(IQR), mmHg | 86.00 (78.00, 93.00) | 88.00 (81.00, 94.00) | 0.285 |
| Smoking, n(%) | 114 (44.71) | 16 (32.65) | 0.118 |
| Drinking history, n(%) | 38 (14.90) | 5 (10.20) | 0.387 |
| Time of onset, Median(IQR), day | 1.20 (0.63,4.00) | 1.25 (0.33,3.00) | 0.423 |
| **Comorbidities, n(%)** |  |  |  |
| Hypertension, n(%) | 200 (78.43) | 37 (75.51) | 0.651 |
| Diabetes mellitus, n(%) | 102 (40.00) | 19 (38.78) | 0.873 |
| Atrial fibrillation, n(%) | 13 (5.10) | 9 (18.37) | 0.003 |
| CHD, n(%) | 32 (12.55) | 11 (22.45) | 0.069 |
| History of stroke, n(%) | 57 (22.35) | 9 (18.37) | 0.535 |
| **Biochemistry, Median(IQR)** |  |  |  |
| WBC, 10<sup>9</sup>/L | 7.00 (5.70, 8.75) | 8.20 (6.60, 9.10) | 0.015 |
| HGB, g/L | 132.00 (121.00, 144.00) | 132.00 (126.00, 141.00) | 0.613 |
| PLT, 10<sup>9</sup>/L | 184.00 (156.00, 224.00) | 194.00 (149.00, 241.00) | 0.822 |
| TC, mmol/L | 4.24 (3.58, 5.06) | 4.30 (3.93, 5.06) | 0.579 |
| TG, mmol/L | 1.59 (1.14, 2.26) | 1.17 (0.94, 1.57) | 0.002 |
| HDL, mmol/L | 1.03 (0.90, 1.19) | 1.14 (0.94, 1.26) | 0.099 |
| LDL, mmol/L | 2.38 (1.85, 3.00) | 2.42 (1.81, 2.88) | 0.964 |
| HbA1c, mmol/L | 6.20 (5.80, 7.75) | 6.10 (5.70, 7.50) | 0.432 |
| CRP, mg/L | 1.00 (0.26, 1.92) | 1.23 (0.50, 8.37) | 0.018 |
| FBG, mmol/L | 5.64 (4.76, 8.10) | 6.03 (5.11, 8.30) | 0.147 |
| CK, U/L | 12.00 (7.00, 15.00) | 10.00 (8.00, 16.00) | 0.674 |
| BUN, mmol/L | 5.56 (4.63, 6.93) | 6.59 (5.49, 7.77) | 0.026 |
| Cr, umol/L | 77.00 (64.00, 94.50) | 75.00 (59.00, 106.00) | 0.992 |
| UA, umol/L | 349.00 (286.50, 408.00) | 347.00 (250.00, 407.00) | 0.476 |
| PT, s | 12.70 (11.80, 13.50) | 13.60 (12.40, 14.20) | 0.001 |
| APTT, s | 30.80 (27.15, 33.65) | 31.70 (29.00, 34.50) | 0.073 |
| TT, s | 16.30 (14.10, 17.15) | 15.90 (14.50, 16.90) | 0.311 |
| FIB, g/L | 3.29 (2.87, 3.79) | 3.52 (2.94, 4.04) | 0.123 |
| PTA, % | 99.70 (91.20, 110.05) | 91.00 (84.00, 102.00) | < 0.001 |
| INR, | 0.99 (0.94, 1.04) | 1.05 (0.98, 1.10) | < 0.001 |
| FT3, pg/mL | 2.94 (2.70, 3.24) | 2.73 (2.31, 3.06) | 0.003 |
| FT, ng/dL | 1.11 (1.01, 1.23) | 1.23 (1.06, 1.33) | 0.005 |
| TSH, mIU/L | 2.24 (1.28, 3.84) | 1.95 (0.94, 2.94) | 0.073 |
| ESR, mm/h | 13.00 (7.00, 26.00) | 20.00 (11.00, 31.00) | 0.114 |
| NIHSS | 2.00 (1.00, 4.00) | 9.00 (6.00, 12.00) | < 0.001 |

IQR: interquartile range; BMI: body mass index; SBP: systolic blood pressure; DBP: diastolic blood pressure; CHD: coronary heart disease; WBC: white blood cell; HGB: hemoglobin; PLT: platelet; TC: total cholesterol; TG: triglyceride; HDL: high density lipoprotein, LDL: low density lipoprotein, CRP: C-reactive protein, FBG: fasting blood glucose, CK: creatine kinase; BUN: blood urea nitrogen; Cr: creatinine, UA: uric acid; PT: prothrombin time; APTT:activated partial thromboplastin time; TT: thrombin time; FIB: fibrinogen,PTA: prothrombin time activity; INR:international normalized ratio; FT3:free triiodothyronine; FT:free thyroxine; TSH:thyroid stimulating hormone; ESR: erythrocyte sedimentation rate; NIHSS: the National Institutes of Health Stroke Scale.

The median (IQR) concentration of TRP was 9877.73 ng/ml (IQR: 8671.87, 11121.88). Within the TRP metabolic pathway, the median concentration of NFK was 5.26 ng/ml (IQR: 3.98, 7.01), KYN was 530.35 ng/ml (IQR: 437.50, 632.81), KYNA was 9.18 ng/ml (IQR: 6.69, 12.27), 3-hydroxykynurenine (3-HKYN) was 5.60 ng/ml (IQR: 3.82, 8.50), and 3-HAA was 3.86 ng/ml (IQR: 2.89, 5.03). In the kynurenine pathway, the median concentration of 5-HTP was 7.37 ng/ml (IQR: 4.67, 9.12), 5-HT was 70.00 ng/ml (IQR: 48.73, 93.62), and 5-HIAA was 20.62 ng/ml (IQR: 15.44, 26.60). In the indole pathway, the median concentration of IAA was 307.76 ng/ml (IQR: 211.97, 451.20).

Figure 1 shows a difference in the distribution of metabolites between the Pgood group and the Ppoor group. The concentration of TRP, NFK/TRP ratio, 5-HTP/TRP ratio, 5-HIAA/TRP ratio, and 3-HAA/3-HKYN ratio were higher in the Pgood group than in the Ppoor group. Conversely, NFK, KYN, KYNA, 3-HKYN, KYN/TRP ratio, KYNA/TRP ratio, 3-HKYN/TRP ratio, 3-HAA/TRP ratio, and KYNA/KYN ratio were higher in the Ppoor group than in the Pgood group.

**Figure 1.**
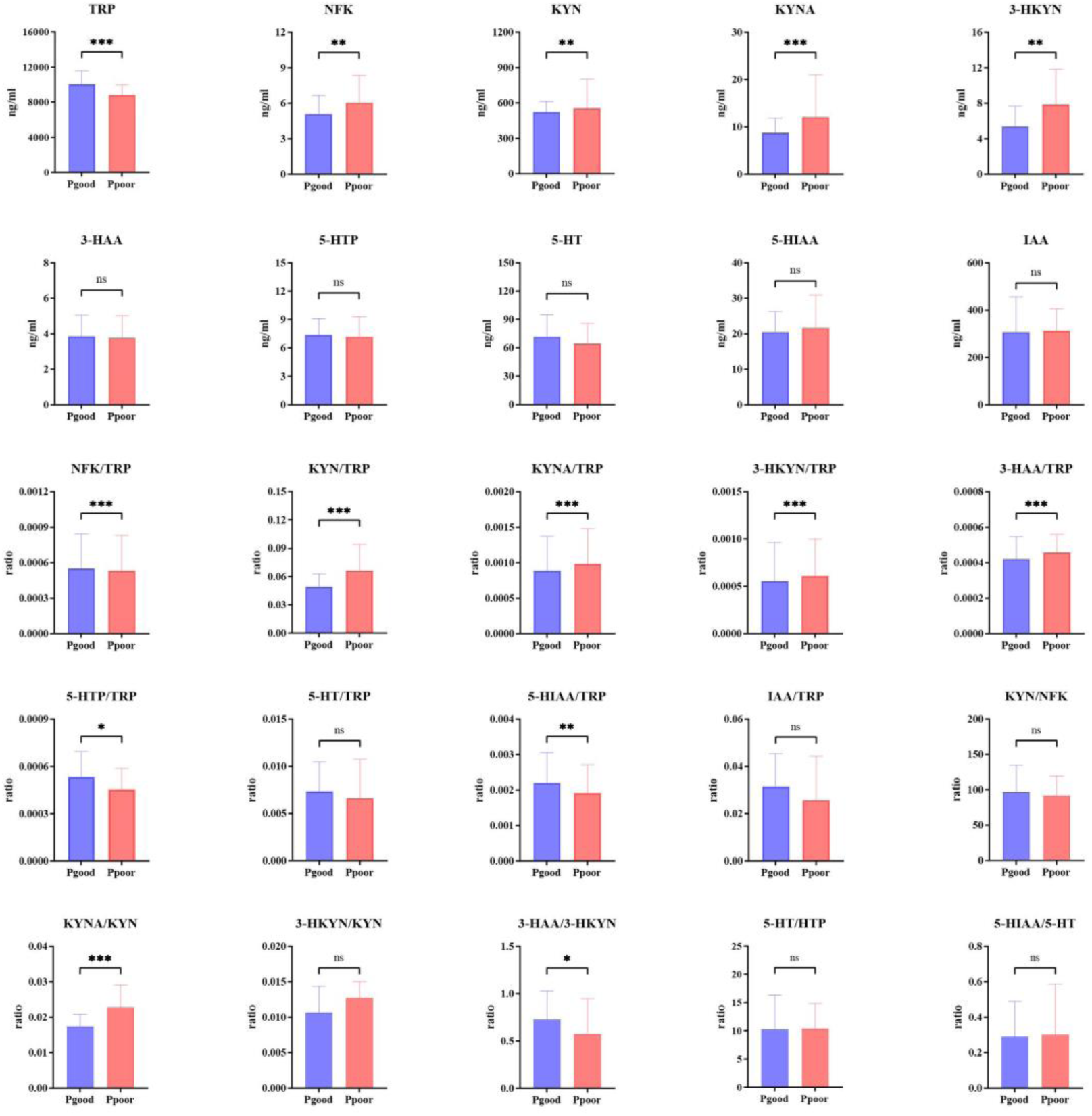
The distribution of tryptophan metabolites and their ratios between the Ppoor and Pgood groups in AIS patients.

### 3.2 Correlations of metabolic parameters

An analysis of the Spearman correlation between 10 metabolites and 15 metabolite ratios in the TRP metabolic pathway, the kynurenine pathway, and the indole pathway was conducted. Except for the significant negative correlations between 5-HT and metabolites of the TRP metabolic pathway, between TRP and 3-HKYN, and between 5-HTP and IAA, all other metabolites showed significant positive correlations with each other (Figure 2A). Moreover, inspection of the correlation matrix revealed mild to moderate correlations among most metabolite ratios (Figure 2B).

**Figure 2.**
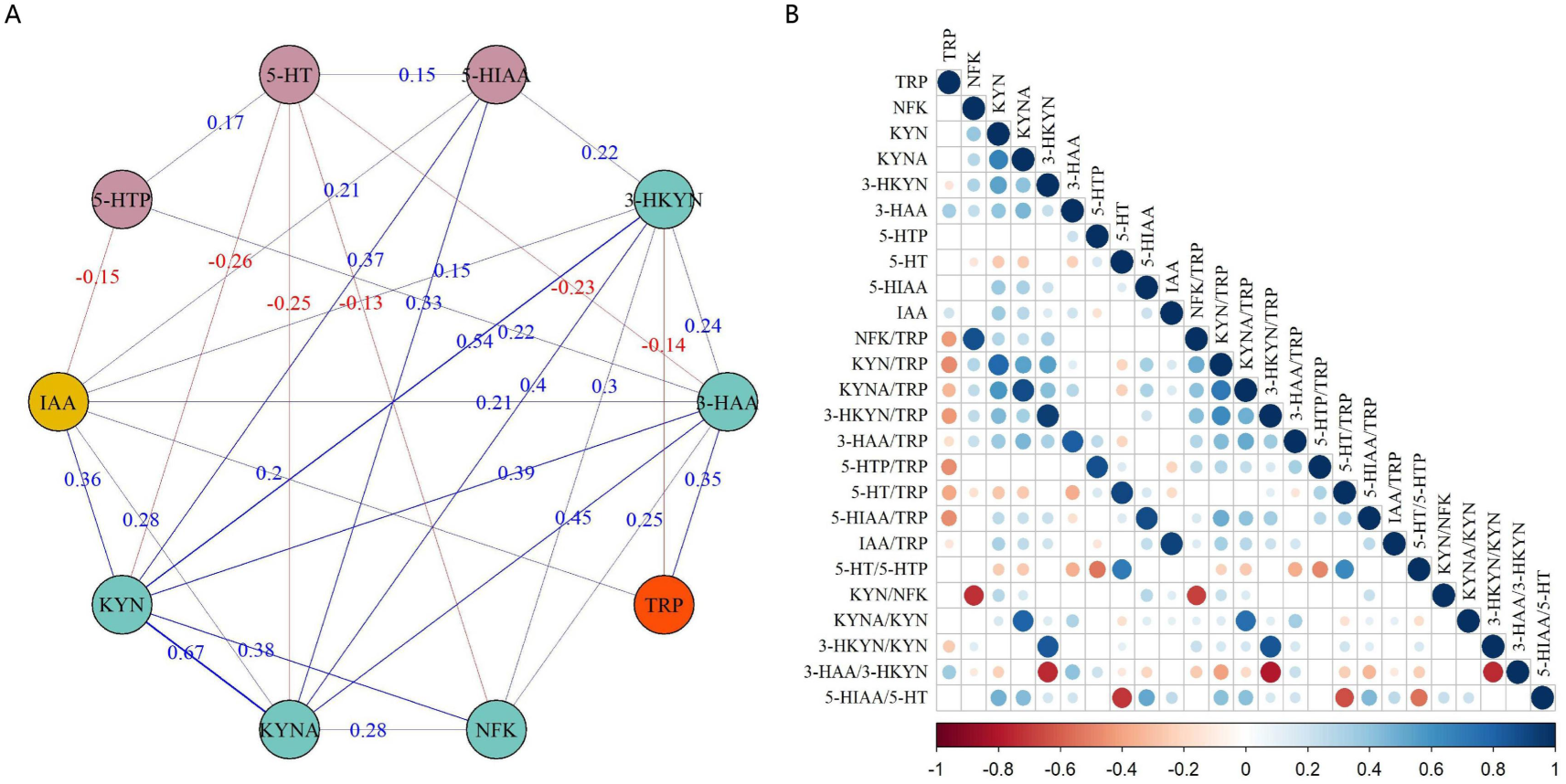
Correlation analysis. (A) Network diagram of correlation between 10 metabolites. Blue lines indicate a positive correlation and red lines indicate a negative correlation. (B) Diagonal matrix plot of 10 metabolites and 15 metabolite ratios. Color represents the correlation coefficient, with red indicating a positive correlation and blue indicating a negative correlation. The size of the circle represents the correlation coefficient.

### 3.3 Correlation of metabolic parameters with disease severity

The lollipop plot was used to visualize the correlation between the NIHSS score and 10 metabolites and their ratios (Figure 3). The NIHSS score exhibited a weak positive correlation with several metabolite parameters, including KYNA/TRP ratio, NFK/TRP ratio, 3-HKYN/TRP ratio, KYNA/KYN ratio, 3-HKYN/KYN ratio, as well as KYNA and 3-HKYN (0 < r < 0.3, *p* < 0.05). In contrast, other metabolite parameters, such as TRP, 5-HTP, and the 3-HAA/3-HKYN ratio, were weakly negatively correlated with the NIHSS score (-0.3 < r < 0, *p* < 0.05). These findings suggest that TRP and its downstream metabolic products may play a role in the severity of AIS.

**Figure 3.**
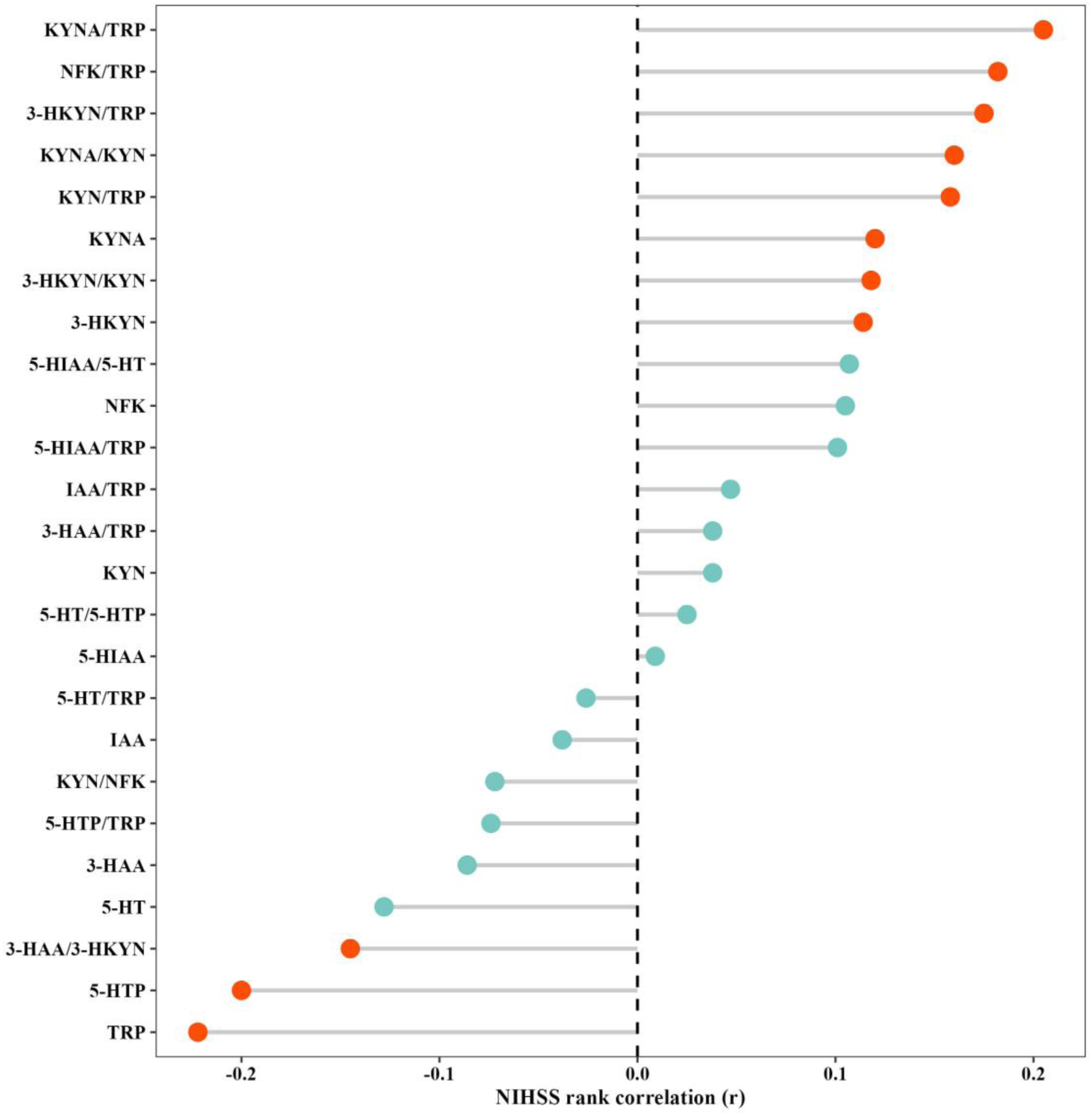
Lollipop plots of correlation between TRP metabolites and NIHSS score. Orange indicates a significant difference (*p* < 0.05), and green indicates no significant difference (*p* ≥ 0.05).

### 3.4 Identification and visualization of cerebral infarction-related potential predictors

The above 12 clinical parameters and 14 metabolite parameters identified through univariate analysis were incorporated into a multivariate analysis to ascertain the independent prognostic factors influencing the 3 months outcome of cerebral infarction. A collinearity analysis demonstrated a VIF > 10, indicating significant collinearity among the independent variables. To ensure the robustness of the results, LASSO regression and random forest models were used to select independent prognostic factors.

LASSO regression analysis was performed to screen out the seven best prognosis-related factors, including TRP, KYN/TRP ratio, KYNA/TRP ratio, age, PTA, NIHSS, and FT4 (Figures 4A, B). According to the constructed optimal risk model, the ROC curve for the prognostic signature is depicted in Figure 4(C). The area under the curve (AUC) was calculated to be 0.917 (95% CI: 0.880, 0.952), with optimal sensitivity and specificity values of 91.8% and 79.2%, respectively. Moreover, a random forest recursive feature elimination approach was employed to identify the minimal set of features essential for differentiating between poor and good prognoses, resulting in the identification of 11 significant features (Figures 4D, E). Figure 4F presents the ranking of feature importance derived from the random forest analysis, highlighting the variables identified as NIHSS, KYNA/TRP ratio, KYN/TRP ratio, age, TRP, KYNA/KYN ratio, KYNA, PT, INR, 3-HAA/3-HKYN ratio, and PTA. The AUC of the optimal risk model of these variables was 1.000 (95% CI: 1.000, 1.000), with optimal sensitivity and specificity values of 100.0% and 100.0%, respectively. The DeLong test showed a statistically significant difference in the AUC under the ROC curve between the LASSO regression and random forest models (*p* < 0.001).

**Figure 4.**
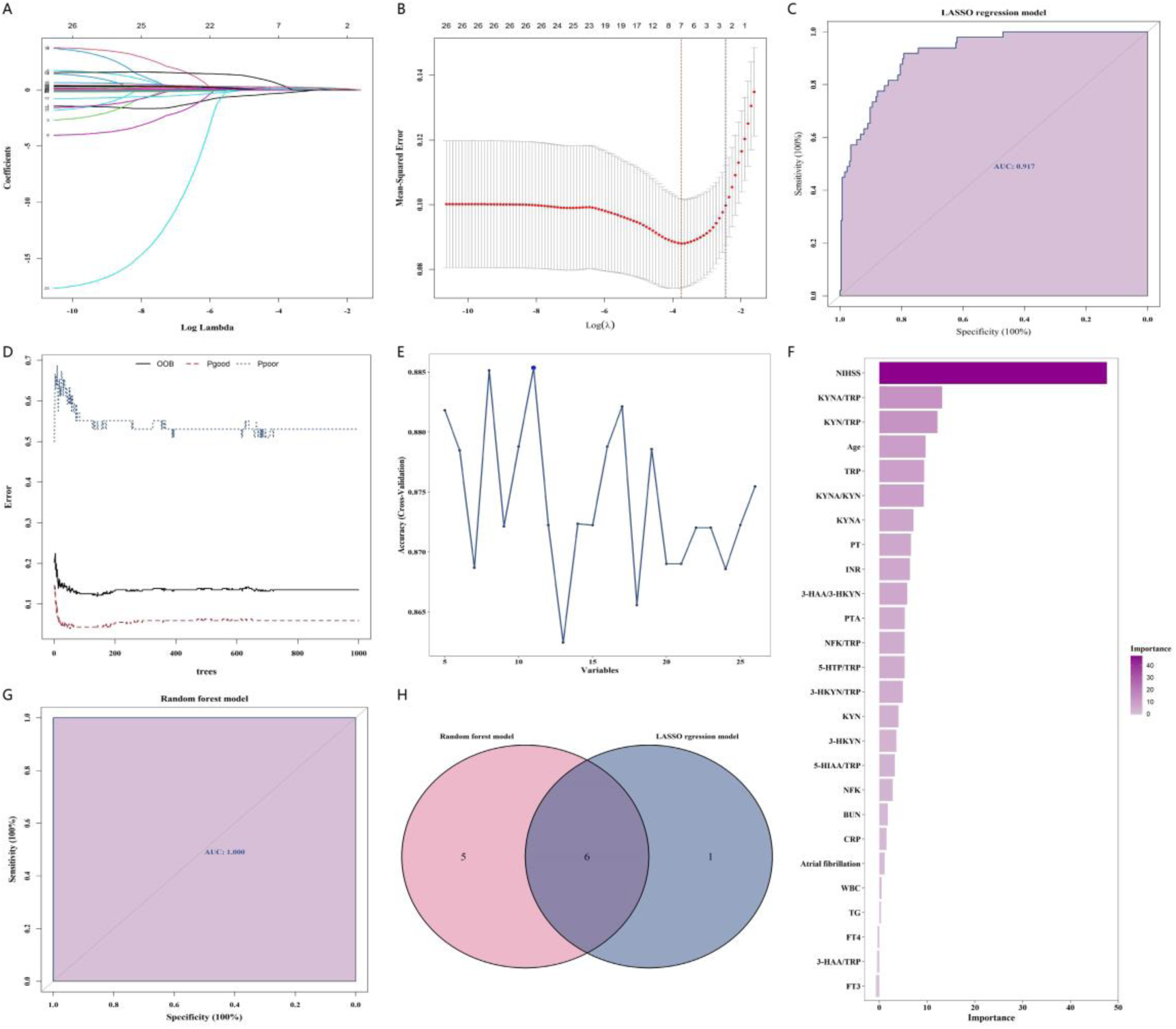
Analysis of LASSO regression and random forest. (A) Plot of the variation of the L1 parity against the regression coefficient. (B) 10-fold CV (cross-validation) MSE (mean squared errors) of lasso regression with varying penalty parameter λ. (C) ROC curve of the LASSO model based on 7 factors. (D) Random forest tree. (E) Automatic feature selection recursive feature elimination with cross-validation. (F) Feature importance rank distribution of random forest model. (G) ROC curve of the random forest model based on 11 factors. (H) Venn diagram of 6 OUTs from both models.

Furthermore, Venn diagram analysis showed that 6 OTUs were shared among the 2 optimal risk models (Figures 4H), including NIHSS, KYNA/TRP ratio, KYN/TRP ratio, TRP, age, and PTA.

### 3.5 Evaluation of the prognostic value of three metabolic parameters

Altogether, one metabolite (TRP) and two ratios of metabolites (KYNA/TRP and KYN/TRP) were determined. Restricted cubic splines were used to assess the dose-response relationship between metabolic parameters and the risk of poor prognosis in cerebral infarction (Figure 5). The analysis revealed a linear dose-response relationship between three parameters of TRP metabolism and the risk of poor prognosis, with the test of nonlinearity not statistically significant (*p* > 0.05).

**Figure 5.**
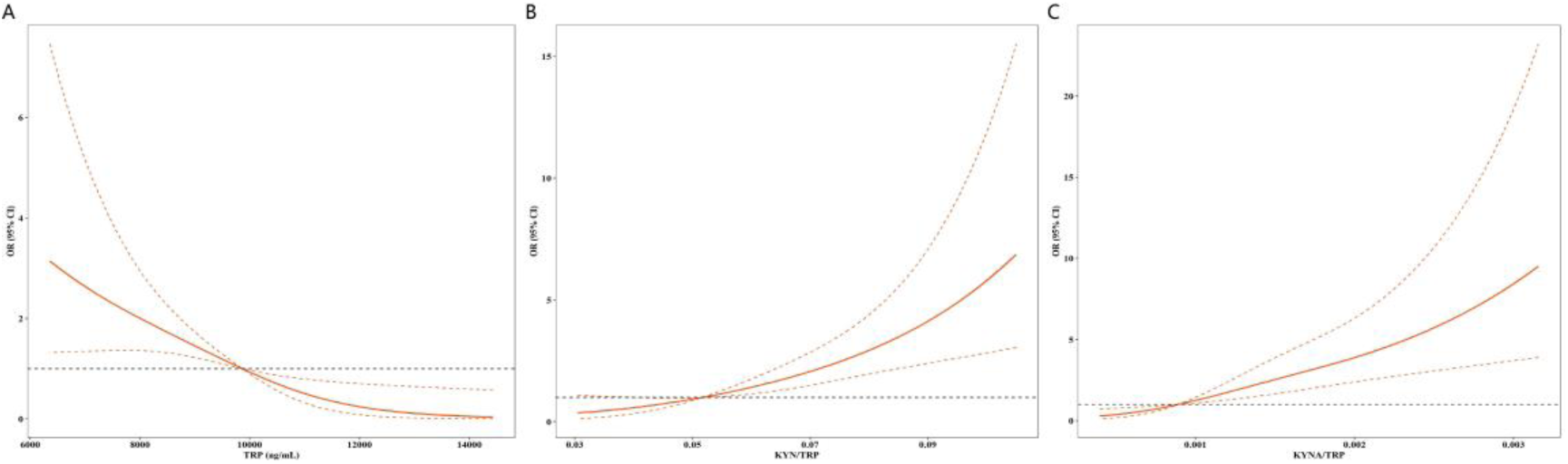
Restricted cubic spline plot of the association between metabolic parameters and the risk of poor prognosis in AIS. The *p*-values for testing the linear relationship among TRP (A), KYN/TRP ratio (B), and KYNA/TRP ratio (C) are 0.286, 0.691, and 0.114, respectively.

Table 2 shows the results of unadjusted (univariable) and adjusted (multivariable) logistic regression analyses. Adjusted for potential confounders, the multivariable logistic regression analysis revealed that both KYN/TRP ratio [odds ration (OR)=2.06, 95% confidence interval (CI): 1.23-3.60, *p*=0.008] and KYNA/TRP ratio (OR=2.15, 95% CI: 1.23-4.12, *p*=0.014) were significantly and positively associated with poor prognosis in cerebral infarction, while TRP was negatively associated with poor prognosis in cerebral infarction (OR=0.46, 95% CI: 0.26-0.76, *p*=0.004). These findings suggest that the OR are robust to adjustments for other relevant confounding variables.

**Table 2.** The association between metabolic parameters and poor prognosis risk in AIS.

| Independent variables | Unadjusted |  | Adjusted |  |
| --- | --- | --- | --- | --- |
|  | OR (95%CI) | *p*-value | OR (95%CI) | *p*-value |
| TRP | 0.40 (0.27, 0.58) | <0.001 | 0.46 (0.26, 0.76) | 0.004 |
| Kyn/TRP ratio | 1.72 (1.29, 3.33) | <0.001 | 2.06 (1.23, 3.60) | 0.008 |
| KYNA/TRP ratio | 2.58 (1.84, 3.78) | <0.001 | 2.15 (1.23, 4.12) | 0.014 |

TRP: OR: odds ratio; CI: confidence interval; TRP: tryptophan; KYN: kynurenine; KYNA: kynurenic acid. Raw data shown after z-score transformation, OR was calculated per one SD increase in biological markers. The adjustment factors included age, atrial fibrillation, WBC, TG, CRP, BUN, PT, PTA, INR, FT3, FT, and NIHSS.

## 4. Discussion

In the previous study, the increased TRP catabolism is initiated before or immediately after stroke, and remained changed in the acute period. In the study of our study, the

Our study revealed that KYN/TRP, KYN/TRP and TRP were associated with stroke severity on hospital admission and poor functional outcome at 3 months even after adjusting for the age, NIHSS score and other factors which are considered strong predictive factors for stroke prognosis. We also developed a model consisting of TRP, KYN/TRP ratio, KYNA/TRP ratio, age, PTA, NIHSS and FT4 to predict the prognosis of cerebral infarction with optimal sensitivity and specificity values of 100.0% and 100.0%, respectively. Our results presented a valuable tool for stroke prognosis predicting and improved the insight in the role of the TRP metabolism in AIS which may contribute to the development of more adequate therapeutics.

The degradation of TRP proceeds via three primary pathways: the KYN pathway, the 5-HT pathway and indole pathway. Within the KYN pathway, there are three key branches involved in its metabolism: the kynurenic acid (KYNA) pathway, the anthranilic acid (AA) pathway and 3-hydroxykynurenine (3-HKYN) pathway [20]. It was reported by Baran *et al.* that an increased synthesis of KYNA and 3-HKYN in the KYN pathway occurred within minutes following a hypoxic episode in neonatal rats, the levels produced showing a direct correlation to mortality[21]. Additionally, KYNA and 3-HKYN could directly interfere with mitochondrial respiration [22], thereby exacerbating the metabolic and oxidative stress in the brain after ischaemia. Darlington et al. demonstrated that the activation of the KYN pathway occurs in AIS patients is with elevated KYN/TRP ratio and decreased 3-HAA/AA ratio which were strongly correlated with infarct volume [23]. Furthermore, a study in 149 AIS patients showed that the KYN/TRP ratio was independently and positively associated with stroke severity [16]. In our study, the KYNA/TRP ratio, NFK/TRP ratio, 3-HKYN/TRP ratio, KYNA/KYN ratio, 3-HKYN/KYN ratio, KYNA and 3-HKYN levels were positively correlated with stroke severity as assessed by NIHSS scores on admission. Conversely, TRP and the 3-HAA/3-HKYN ratio were negatively correlated with NIHSS scores. These findings suggest the activation of the KYN and 3-HKYN pathways in AIS patients. Additionally, we demonstrated that 5-HTP, a key component in the 5-HT pathway was negatively correlated with NIHSS. It was shown that the decrease in 5-HTP CSF levels correlated with frontal atrophy which might reflect the time-dependent changes in phenylketonuria patients [24].

Our study found that in AIS patients who had a poor prognosis 3 months later, the KYN and KYNA pathway of TRP metabolism were activated, as evidenced by increased KYN/TRP and KYNA/TRP ratios along with decreased TRP levels.The elevated KYN/TRP ratio suggests increased activity of indoleamine 2,3-dioxygenase (IDO). Proinflammatory cytokines induce the expression of IDO which catalyzes the rate-limiting step in the production of KYN from TRP. An elevated KYN/TRP has been associated with inflammatory and neurodegenerative conditions and with clinically important outcomes, including mortality in the very old [25], poorer cognitive function in those with Alzheimer’s disease [26] or the severity of bacteremia patients [27]. It was proved in the study of Mo et al. that KYN/TRP was positively associated with the NIHSS at 3 weeks post-stroke [28]. However, this study had certain limitations, including a small sample size (n = 81) of patients with AIS and without adjusting for multiple factors. Our study revealed the prognostic value of KYN/TRP after adjusting for NIHSS, age and other risk indicators in a larger cohorts.

An increased KYNA/TRP indicates the activation of KYNA pathway and the increased activity of kynurenine aminotransferase. It is evident that maintaining an appropriate balance of KYNA is crucial [29], as elevated levels have been implicated in patients with herpes simplex virus type 1 encephalitis [30], while decreased levels have been associated with major depression [31] and Parkinson’s disease [32]. Increased KYNA concentrations will block glutamate receptors in the brain [33], and might represent an adaptive response to block the potentially harmful effects of excessive glutamate receptor stimulation following ischaemia. This blockade of a major excitatory neurotransmitter could further depress brain function, especially in severely affected stroke patients, leading to the increased mortality [23]. It might be presumed that the elevated levels of KYNA/TRP associated with poor prognosis are a result of the body’s failed attempt at protecting cells. Decreased TRP levels in the blood, a consequence of the activation of KYN pathway, limit the transport of the amino acid across the blood-brain-barrier and diminish 5-HT availability in the brain. This reduction may manifest as cognitive decline, depressive symptoms [34] and reduced rehabilitation willingness.

Identification of the KYN/TRP, KYN/TRP and TRP as biomarkers may represent a significant step toward improving long-term stroke prognosis, as kynurenine metabolites are potentially amenable to pharmacological manipulation[33]. In the central nervous system, kynurenine aminotransferase II, predominantly expressed in astrocytes, is responsible for KYNA synthesis, whereas kynurenine 3-monooxygenase (KMO), primarily localized in microglia, catalyzes the production of 3-HYKN, which subsequently leads to quinolinic acid formation [35]. In animal models of cerebral ischemia, administration of kynurenine sulfate exerts neuroprotective effects and notably increases cerebral concentrations of KYNA [36]. It is now recognized that cerebral concentrations of 3-HKYN and quinolinic acid can be selectively manipulated without altering KYNA levels through inhibition of KMO [35], an approach that demonstrates neuroprotective effects in animal models of cerebral ischemia [37].

In previous studies investigating TRP metabolism in AIS patients, the primary focus was on the KYN pathway [16]. However, the products in the 5-HT and indole pathway are also implicated in the AIS. For instance, 5-HT plays a crucial role in neurotransmission, vasoconstriction, or vasodilation of blood vessels, control of hemostasis and platelet function [38]. Melatonin is also involved in the interactions between the nervous, endocrine, and immune systems and serves as a beneficial agent in the treatment of inflammatory and immune disorders [39, 40] including cerebral vascular disease. In this study, we examined the metabolites resulting from three TRP degeneration pathways. Our analysis did not reveal any significant correlations between the prognosis of AIS and metabolites from the 5-HT or indole pathways, which might be attributed to the existence of the blood-brain barrier.

As TRP is an essential amino acid, it must be obtained through diet. Therefore, it is conceivable that TRP concentrations may vary between different dietary habits and races[41]. To our knowledge, there have been few studies on the correlation between the long term prognosis of AIS and TRP in Chinese mainland population.

This study had some limitations. First, it was completed at a single center, therefore, further research is required using multicenter prospective cohorts. Second, the TRP and catabolites were measured at admission, and a repeated-measures analysis may have captured a larger proportion of the TRP and catabolites variance. However, this early time point is the most useful for identifying a possible biomarker-based risk stratification. Third, there is a possibility of connection between dietary TRP consumption [42]. However, we had no information about dietary habits of the patients. Besides, it would be interesting to measure the concentrations of enzyme in the TRP metabolism.

## 5. Conclusion

This study investigated the extraction of three primary variables — TRP, KYN/TRP ratio, and KYN/TRP ratio—from tryptophan metabolic features utilizing LASSO and random forest algorithms to predict the 3-month prognosis of patients with AIS. The findings suggest some new potential prognostic biomarkers or molecular targets for patients with AIS.

## Author Contributions

Conception and design, C.P, C.Q;Performed the experiment, F.C and H.L; Data analyses, Y.Y and C.Q; Manuscript preparation and Manuscript writing, C.P, Y.Y and C.Q; Reviewing and Critical revision, C.Q.

## Acknowledgments

We expressed our gratitude to all participants and staff involved in the study.

## Data Available Statement

The data utilized to bolster the results of this research are incorporated within the article.

## Institutional Review Board Statement

This study was conducted in accordance with the Declaration of Helsinki, and approved by the Ethics Committee of The Hunan University of Medicine General Hospital.

